# Multiple pathways prevent bi-parental mitochondria transmission in *C. elegans*

**DOI:** 10.1101/2025.03.18.642775

**Authors:** Valentine Melin, Justine Cailloce, Fanny Husson, Jorge Merlet, Vincent Galy

## Abstract

The uniparental transmission of the maternal mitochondrial genome is achieved by the disposal of the sperm mitochondria and their DNA (mtDNA) around fertilization. *C. elegans* embryo allowed the identification of several factors and mechanisms involved after fertilization. The discovery that the autophagy pathway and its adaptor ALLO-1 are implicated represented a major progress. In addition, studies in fish and fly have implicated CPS-6, a sperm mitochondrial endonuclease, in the degradation of *C. elegans* sperm-derived mtDNA. The sperm mitochondria-associated proteins FNDC-1 and PHB-2 were also implicated in efficiently removing sperm-derived mitochondria. Despite these recent findings, it is still not clear whether they account for the complete degradation and if we can experimentally prevent sperm-mitochondria clearance and induce sperm-mtDNA transmission to the progeny.

Here, we investigated the genetic interactions and tested the impact of simultaneous inactivation of known factors on the fate of sperm-derived mitochondria and their mtDNA. We revealed an additive effect of ALLO-1 and CPS-6 loss of function. Furthermore, the inactivation of known factors acting in sperm mitochondria clearance was not sufficient to avoid sperm mitochondria removal and to allow paternal mtDNA inheritance across generations.

These findings reveal that *C. elegans* employs at least two parallel pathways to ensure the degradation of paternal mitochondria and suggest the presence of unidentified mechanism(s) safeguarding maternal mitochondrial inheritance.

## INTRODUCTION

Mitochondria are essential organelles that produce adenosine triphosphate (ATP) and control many cellular processes. Mitochondria have their genome, the mitochondrial DNA (mtDNA). In most eukaryotes, contrary to the nuclear genome which is inherited equally from both parents during sexual reproduction, the mtDNA is only inherited from the mother. It is now well established that maternal mitochondrial heredity is not only the result of dilution of the sperm-derived mtDNA over the maternal mtDNA. Indeed, it relies on various active mechanisms that prevent the transmission of the sperm mitochondrial genome (Birky, 2001). Even though uniparental maternal heredity is a conserved phenomenon, the elimination of sperm mitochondrial genome occurs at distinct stages of the reproductive cycle, depending on the species.

In *C. elegans*, as in most animal species, the sperm mitochondria and their genomes enter the oocyte upon fertilization. But within 20-25 minutes after fertilization, a strong and selective autophagy wave named allophagy is induced against sperm-derived mitochondria (Al Rawi et al., 2011; Sato and Sato, 2011). The maternal autophagy receptor/adaptor ALLO-1 is, to date, the earliest factor recruited around the two derived sperm organelles aimed to be eliminated in worms: the sperm mitochondria and the membranous organelles (MOs), a Golgi-derived sperm organelles (Al Rawi et al., 2011; Sato et al., 2018; Sato and Sato, 2011).

Several other conserved factors were described to participate in sperm-derived mitochondria degradation (Merlet et al., 2019). Among them, the mitochondrial endonuclease G (CPS-6 in *C. elegans*) acts as a paternal factor required for sperm mitochondria and mtDNA degradation (Zhou et al., 2016). Unlike in *C. elegans* where CPS-6 acts after fertilization, in *Drosophila*, endonuclease G contributes to uniparental maternal mitochondrial inheritance before fertilization during the sperm individualization process. Two other paternal conserved factors, known for their role in mitophagy in somatic cells, also participate in sperm-derived mitochondria degradation in *C. elegans* after fertilization : FNDC-1 - the ortholog of human FUNDC1 - and PHB-2 a mitochondrial inner membrane receptor (Lim et al., 2019; Wei et al., 2017). A single inactivation of one of these four factors does not allow for a stable transmission to the following generations. In addition, little is known about their genetic interaction(s) and if they can account for the complete mechanisms sustaining uniparental maternal inheritance or if additional mechanism(s) exist.

To better understand the genetic interaction(s) between ALLO-1, CPS-6, FNDC-1, and PHB-2 in sperm-derived mitochondria degradation, we simultaneously inactivated these factors. We revealed an additive effect of ALLO-1 and CPS-6 loss of function indicating that they act in parallel pathways. However, the simultaneous inactivation of those four factors did not allow sperm mitochondrial inheritance. Therefore, our results suggest that in *C. elegans*, at least two pathways act in parallel to ensure the proper degradation of sperm-derived mitochondria. Our results also suggest that additional mechanism(s), which remain(s) to be identified, ensure mtDNA removal and prevent transmission to the next generation.

## RESULTS

### Simultaneous inactivation of *cps-6* and *allo-1* showed a synergistic effect on sperm mitochondria stabilization

To establish the respective contribution and potential functional interaction between *allo-1* and *cps-6*, we tested if depleting both factors impacted on the stabilization of sperm-derived mitochondria. In the *cps-6(tm3222)* allele, the deletion removes the catalytic site of CPS-6 which is required for endonuclease activity and to mediate sperm-derived mitochondria elimination (Parrish et al., 2001; Zhou et al., 2016). In the *allo-1(tm4756)* allele, the deletion creates a premature stop codon which results in a truncated protein without its LC3-interacting region (LIR) motif, required for LGG-1 recruitment - the mammalian homolog of LC3/Atg-8 (Sato et al., 2018). For each different genetic background tested, males expressing HSP-6::GFP in the germline were crossed with hermaphrodites of the corresponding genotype. In order to prevent auto-fertilization and analysis of embryos from un-labeled spermatozoa, we introduced the feminizing *fem-2(b245)* mutation that prevents sperm formation when worms develop at 25°C (Kimble et al., 1984). We crossed feminized worms with males expressing a mitochondrial HSP-6::GFP and recovered the embryos. To overcome the variability associated with counting fluorescently labeled sperm-derived mitochondria, we developed a semi-autonomous method to quantify sperm-derived mitochondria in *C. elegans* embryos (***Figure 1A***).

**Figure 1.**
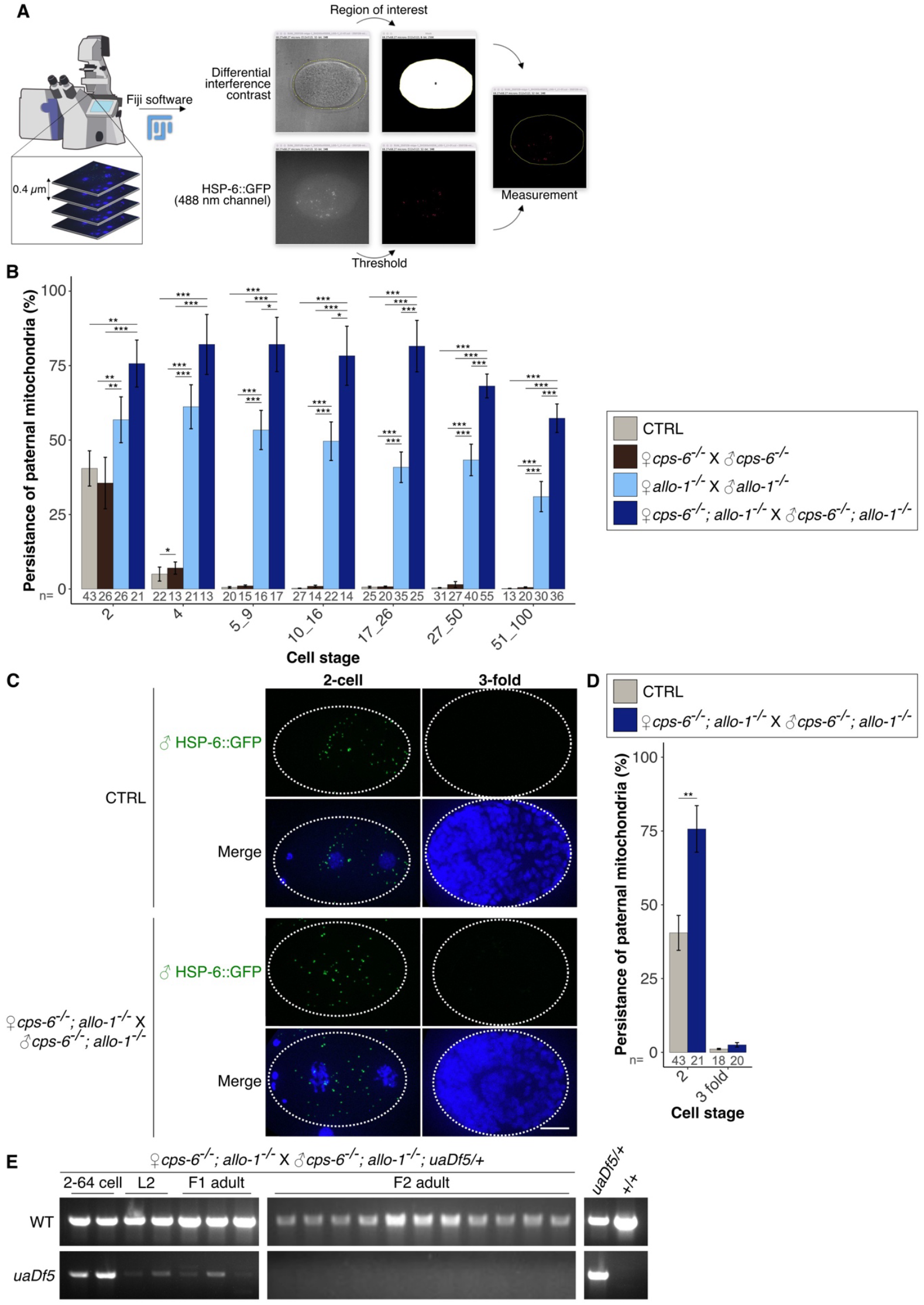
Simultaneous inactivation of *cps-6* and *allo-1* showed a synergistic effect on sperm mitochondria stabilization. **(A)** After orthogonal projection of the z-stacks from embryos imaged with a confocal microscope, the region of interest corresponding to the embryo was delimited and a threshold applied to remove background noise. The relative amount of sperm mitochondria per embryo corresponds to the sum of fluorescence intensity from each stack. **(B)** Quantification of the HSP-6::GFP labeled sperm-derived mitochondria (as % of the 1-cell stage) from the 2-cell stage to 100-cell stage. **(C)** Maximum intensity projections of confocal images of HSP-6::GFP sperm-derived mitochondria (green) and nuclear DNA (blue) in 2-cell and late stage (3-fold) in control and *cps-6(tm3222); allo-1(tm4756)* double mutant and **(D)** quantification of HSP-6::GFP labeled sperm mitochondria (as % of the 1-cell stage). **(E)** PCR amplification of paternally transmitted mitochondrial DNA (*uaDf5*) and wild-type (+) mitochondrial haplotypes in 2 to 64-cell stage embryos, L2 and adults offspring and in F2 adults in the *cps-6(tm3222); allo-1(tm4756)* double mutant. Wilcoxon test p-values: *p<0.05; **p<0.01; ***p<0.001. Scale bar: 10µm.

In *cps-6(tm3222)* embryos, the average HSP-6::GFP intensity for the remaining sperm-inherited mitochondria at the 4-cell stage was slightly different from control embryos (7.0% ± 2.0% and 5.0% ± 2.4% respectively). Whereas, as expected, *allo-1(tm4756)* embryos displayed a stabilization of sperm mitochondria with a remaining signal intensity of 61.2% ± 7.4% at 4-cell stage. Noticeably, double mutant embryos exhibited a significantly higher persistence of sperm mitochondria signal when compared to the single *allo-1(tm4756)*, with a signal intensity of 82.1% ± 10.1% at 4-cell stage (***Figure 1B***). The stabilizing effect in the double mutant was still visible at the 51-100-cell stage. Indeed, only 0.2% ± 0.1% of the signal intensity remained in the control, compared to 31.0% ± 5.1% in *allo-1(tm4756)* and 57.3% ± 4.8% in double mutant embryos. Therefore, our results revealed a synergistic effect of the simultaneous inactivation of *allo-1* and *cps-6* (***Figure 1B***).

We then analyzed if sperm-derived mitochondria were stabilized in older embryos. At the 3-fold stage, the quantification data showed no difference between control and double mutant embryos (***Figure 1C and 1D***). We then analyzed by PCR how long sperm mtDNA was maintained in double mutants. As expected from our quantifications, the sperm-contributed mtDNA, *uaDf5* allele, was amplified in F1 early embryos. To better evaluate the possibility that sperm mtDNA could be maintained in F1, we looked for the *uaDf5* allele on F1 single adults. We found that 27/43 adults F1 contained the *uaDf5* allele. However, none of the F2 progeny from *uaDf5* positive F1 contained the *uaDf5* allele (***Figure 1E***).

We then evaluated whether *cps-6(tm3222)* and *allo-1(tm4756)* alleles cause deleterious defects. We measured the brood size of the adult hermaphrodites N2 (control), single, and double mutants. We observed a modest but significant reduction of the brood size in each single mutants (245 ± 50 for *cps-6(tm3222)*, 248 ± 56 for *allo-1(tm4756)*) as well as in the double mutant (215 ± 52 for *cps-6(tm3222); allo-1(tm4756)*) compared to control (290 ± 47 for N2) (***Figure S1A***). This modest decrease in the brood size was not combined with a reduction in embryonic viability since we did not observe any significant reduction in the fraction of hatching embryos in none of the mutants’ backgrounds (0.5% ± 0.5% for *allo-1(tm4756)*, 1.1% ± 1.6% for *cps-6(tm3222)* and 0.6% ± 1.0% for *cps-6(tm3222); allo-1(tm4756)*) compared to control (0.4% ± 0.4% for N2) (***Figure S1B***).

Taken together, our results showed that, despite a greater persistence of sperm mitochondria in the embryos in double mutant *cps-6(tm3222); allo-1(tm4756)*, this is not enough to maintain the paternal mtDNA over the generations.

### The combined inactivation of ALLO-1, CPS-6, FNDC-1 and PHB-2 is not sufficient to transmit paternal mtDNA over the generations

The simultaneous inactivation of ALLO-1 and CPS-6 was not sufficient to stabilize sperm mitochondria or their mtDNA across the generations. Therefore, we tested the effect of simultaneously inactivating ALLO-1 and CPS-6 endonuclease, along with the depletion of two other known sperm factors FNDC-1 and PHB-2. In order to achieve this goal and avoid potential deleterious effects on development, we crossed feminized hermaphrodites containing *allo-1(tm4756)* mutation with males mutants *cps-6(tm3222); fndc-1(rny14); phb-2(RNAi)* expressing HSP-6::GFP. Feminized worms were also treated with *allo-1(RNAi)* to avoid the embryonic expression of the paternally *allo-1* contributed gene.

We measured the stability of the sperm-inherited mitochondria as previously described. The depletion of FNDC-1 and PHB-2 did not increase the stability of sperm-derived mitochondria compared to the double *cps-6(tm3222); allo-1(tm4756)* mutant embryos neither at early nor late stages (***Figure 2A, 2B and 2D***). Even though sperm-derived mitochondria are visible in some embryos, the remaining GFP signal associated with sperm-derived mitochondria at 3-fold stage was not significantly different in the quadruple mutant than in the double mutant (3.0% ± 0.8% and 2.5% ± 0.8% for *cps-6(tm3222); fndc-1(rny14); phb-2(RNAi); allo-1(tm4756)* and *cps-6(tm3222); allo-1(tm4756)* mutants respectively). Our results showed that depletion of ALLO-1, CPS-6, FNDC-1 and PHB-2 does not differ from the double *cps-6(tm3222); allo-1(tm4756)* mutant, suggesting that sperm-derived mitochondria are still degraded in the absence of functional ALLO-1, CPS-6, FNDC-1 and PHB-2. We obtained similar results by tracking sperm-derived mitochondria with CMXRos, a fluorescent cationic mitochondrial marker (***Figure 2C***). To analyze sperm mtDNA persistence, we followed by PCR the sperm provided *uaDf5* allele. We were able to detect the *uaDf5* allele in 17/36 adults F1 (***Figure 2E***). However, inactivation of these four factors does not allow amplification in none of the F2 animals tested from an *uaDf5* positive L1 adult.

**Figure 2.**
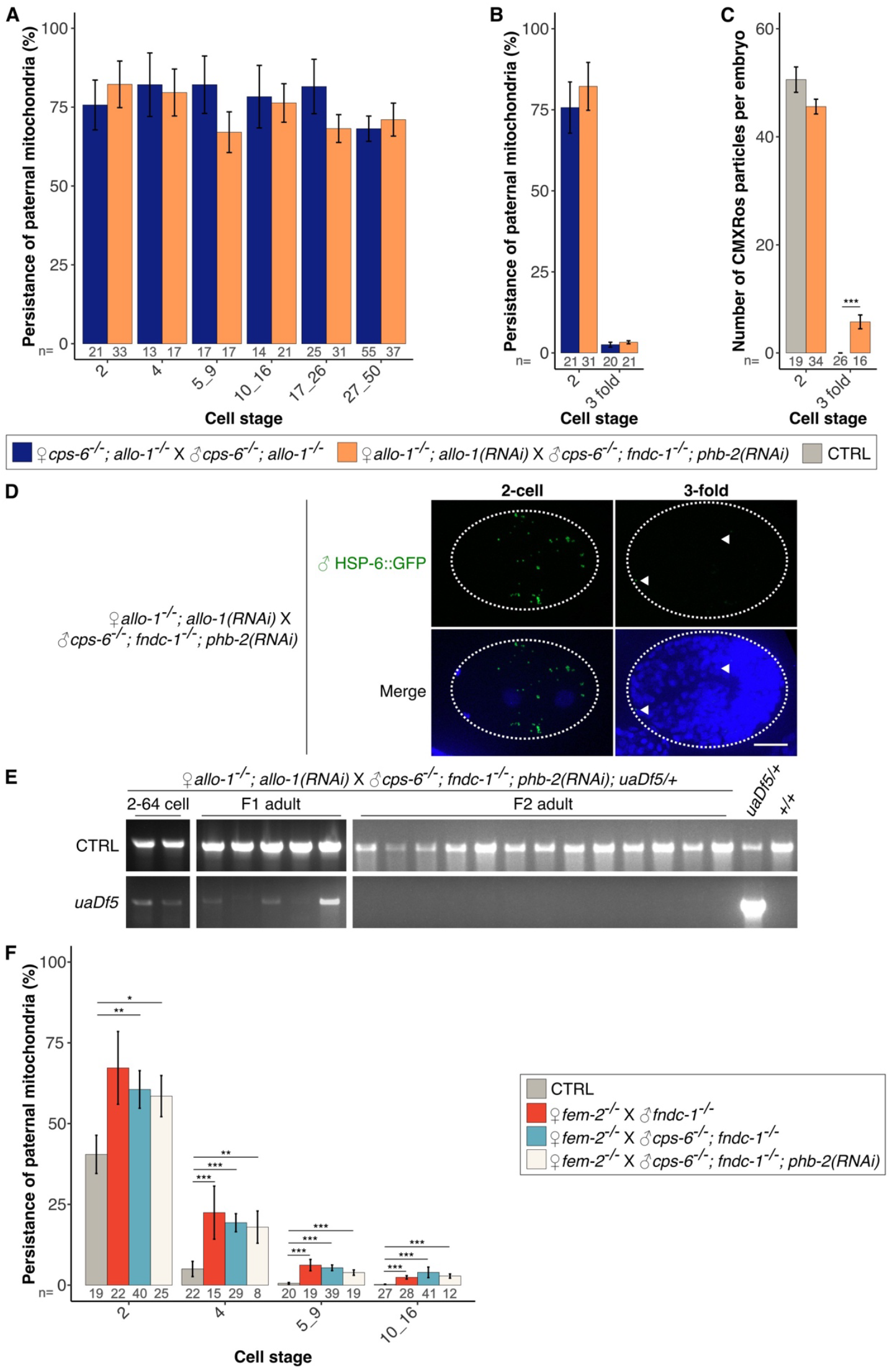
Inactivation of maternal and paternal specific factors did not prevent sperm-derived mitochondria degradation. **(A)** Quantification of the HSP-6::GFP labeled sperm-derived mitochondria (as % of the 1-cell stage) from the 2-cell to 100-cell stage in the *cps-6(tm3222); allo-1(tm4756)* and *cps-6(tm3222); fndc-1(rny14); phb-2(RNAi); allo-1(tm4756)* mutants. **(B)** Quantification of the HSP-6::GFP labeled sperm-derived mitochondria (as % of the 1-cell stage) at the 2-cell and 3-fold stage in *cps-6(tm3222); allo-1(tm4756)* and *cps-6(tm3222); fndc-1(rny14); phb-2(RNAi); allo-1(tm4756)* mutants. **(C)** Quantification of sperm mitochondria labeled with CMXRos in the embryos from the indicated crosses. **(D)** Maximum intensity projections of confocal images of HSP-6::GFP sperm-derived mitochondria (green) and nuclear DNA (blue) in 2-cell and late stage (3-fold) in *cps-6(tm3222); fndc-1(rny14); phb-2(RNAi); allo-1(tm4756)* mutant. **(E)** PCR amplification of paternally transmitted (*uaDf5*) and wild-type (+) mtDNA haplotypes in 2-64-cell stage embryos, L2, adults offspring and in F2 adults from the indicated crosses. **(F)** Quantification of the HSP-6::GFP labeled sperm-derived mitochondria (as % of the 1-cell stage) from the 2-cell to 16-cell stage from crosses with feminized adults with males *fndc-1(rny14)*, or *cps-6(tm3222); fndc-1(rny14)* or *cps-6(tm3222); fndc-1(rny14); phb-2(RNAi)* mutants. Wilcoxon test p-values: *p<0.05; **p<0.01; ***p<0.001. Scale bar: 10µm.

Our results from the double *cps-6(tm3222); allo-1(tm4756)* and quadruple *cps-6(tm3222); fndc-1(rny14); phb-2(RNAi); allo-1(tm4756)* mutants suggested that FNDC-1 and PHB-2 could act either in ALLO-1 pathway or CPS-6 pathway. ALLO-1, FNDC-1 and PHB-2 ability to directly induce autophagosomes formation through their LIR domain suggests that they may act in parallel pathways. Therefore, we tested whether FNDC-1 and PHB-2 could act together or not with CPS-6. We measured the persistence of sperm mitochondria as previously described in embryos from crosses between feminized hermaphrodites and males mutants for *fndc-1(rny14)* or *cps-6(tm3222); fndc-1(rny14)* or *cps-6(tm3222); fndc-1(rny14); phb-2(RNAi)* expressing HSP-6::GFP in the germline. At each stage of embryonic development, no significant differences in GFP signal were observed between single *fndc-1(rny14)*, double *cps-6(tm3222); fndc-1(rny14)* and triple *cps-6(tm3222); fndc-1(rny14); phb-2(RNAi)* mutants suggesting that CPS-6, FNDC-1 and PHB-2 act in the same genetic pathway (***Figure 2F***).

### Sperm-derived mitochondria are not poly-ubiquitylated in late embryos

The combined inactivation of known factors involved in sperm-derived mitochondria degradation induced a stabilization without preventing their clearance at later stages of development (***Figure 2***). To elucidate a potential involvement of poly-ubiquitylation in the targeting of transiently stabilized sperm-derived organelles for degradation, we tested the presence of poly-ubiquitin chains using Tandem-repeated Ubiquitin-Binding Entities (TUBEs).

We measured the stability of sperm-derived mitochondria and MOs in *cps-6(tm3222); fndc-1(rny14); phb-2(RNAi); allo-1(tm4756)* quadruple mutant embryos at late stages, as well as the percentage of these organelles labeled or not with Pan-TUBEs which recognize multiple types of poly-ubiquitin chains, including K48 and K63 chains. As expected from our previous results (Cailloce et al., 2023a), MOs are predominantly poly-ubiquitylated at the 2-cell stage in both control and mutant embryos (66.5% ± 20.8% and 95.5% ± 14.5%, respectively). At later developmental stages in *cps-6(tm3222); fndc-1(rny14); phb-2(RNAi); allo-1(tm4756)* mutant embryos, MOs remained poly-ubiquitylated with 76.6% ± 17.8% of MOs positively labeled at the bean stage and 70.8% ± 34.3% at the 1.5-fold stage (***Figure 3A* and *3C***). However, in late mutant embryos, TUBEs signal remained not detectable in sperm-derived mitochondria with no labeling at bean and 1.5-fold stages (***Figure 3B* and *3D***) like in early embryos (Cailloce et al., 2023a). These results suggested that, although MOs are predominantly poly-ubiquitylated, sperm-derived mitochondria do not exhibit a positive signal for Pan-TUBEs labeling, even when stabilized in embryos.

**Figure 3.**
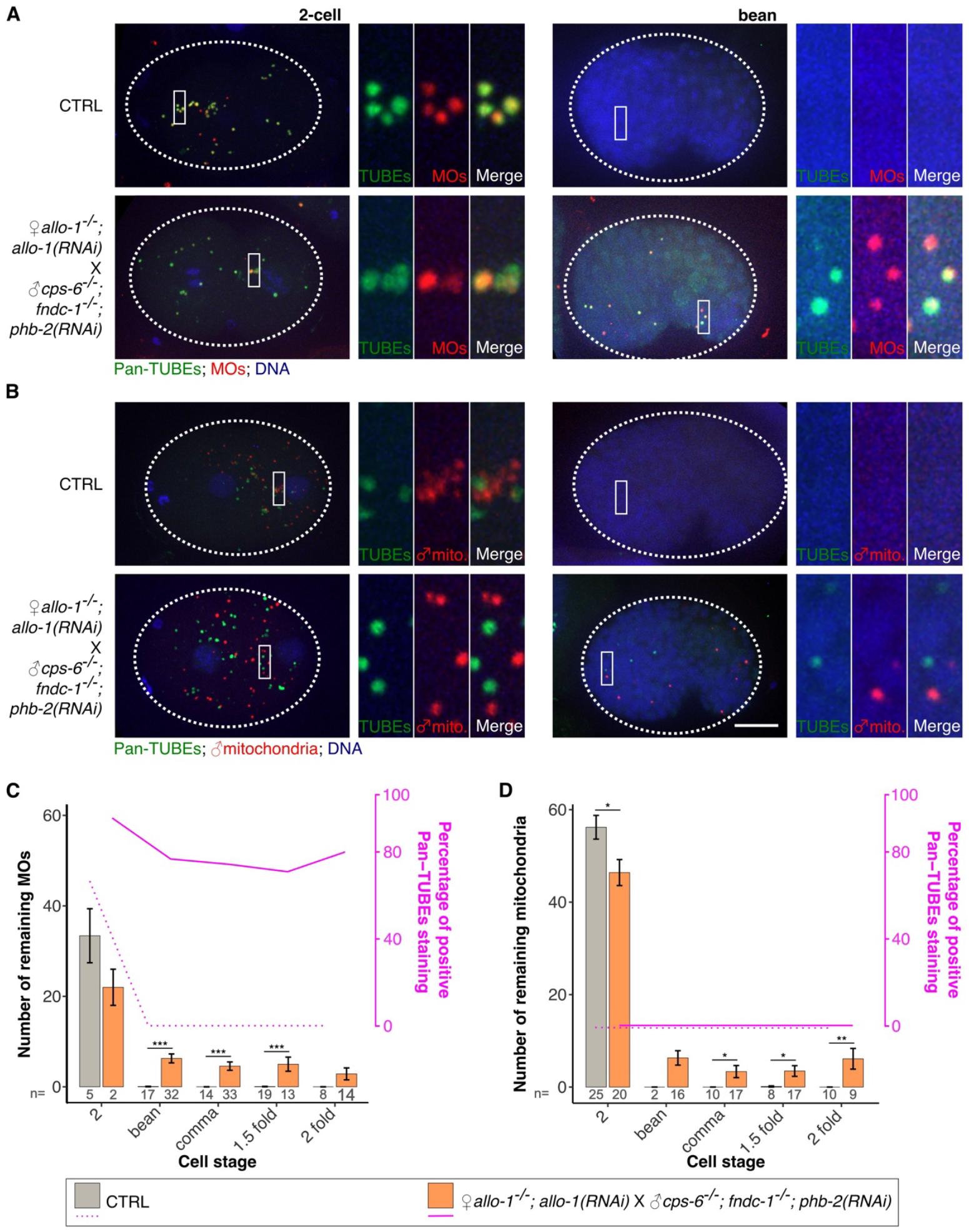
No poly-ubiquitylation detected on sperm-derived mitochondria in control nor in the absence of ALLO-1, CPS-6, PHB-2 and FNDC-1. **(A, B)** Maximum intensity projections of representative control and *cps-6(tm3222); fndc-1(rny14); phb-2(RNAi); allo-1(tm4756)* mutant early and bean stage embryos labeled with Pan-TUBEs (green) and for **(A)** MOs (red) and **(B)** sperm-derived mitochondria (red). **(C)** Quantification of the percentage of MOs and **(D)** sperm-derived mitochondria labeled with Pan-TUBEs in control and *cps-6(tm3222); fndc-1(rny14); phb-2(RNAi); allo-1(tm4756)* mutants embryos at the indicated developmental stages. Wilcoxon test p-values: *p<0.05; **p<0.01; ***p<0.001. Scale bar : 10µm.

### PINK-1 can target sperm-derived mitochondria in *allo-1(tm4756)* mutant embryos

To further evaluate a role of poly-ubiquitylation in sperm-derived mitochondria degradation, we tested if in *allo-1(tm4756)* background the PINK1/Parkin mitophagy pathway could take over the ALLO-1 pathway. We measured sperm-derived mitochondria stability in the *allo-1(tm4756)* embryos upon *pink-1* inactivation. This was achieved by crossing *pink-1(ok3538); allo-1(tm4756)* mutant hermaphrodites, treated by *pink-1* RNAi to avoid embryonic expression of the paternally contributed gene, with *allo-1(tm4756)* males expressing HSP-6::GFP. We measured the remaining sperm-derived mitochondria at different stages of development and established the kinetic of degradation for *allo-1(tm4756)* and *pink-1(ok3538); allo-1(tm4756)* mutants. In agreement with published results showing that the PINK1/Parkin mitophagy pathway was not involved in sperm-derived mitochondria degradation in *C. elegans* (Sato et al., 2018), we did not observe any difference in GFP intensity in early embryos (data not shown) and late embryos (3-fold stage) between single *allo-1(tm4756)* and double *pink-1(ok3538); allo-1(tm4756)* mutants (***Figure 4***). Unexpectedly, we observed a significant but limited increase of the GFP signal between the 10 and 100-cell stages. The increase in sperm-derived mitochondria signal was of 77.3% ± 5.4% compared to 49.6% ± 6.5% respectively in *pink-1(ok3538); allo-1(tm4756)* double mutant and in *allo-1(tm4756)* at 10-16-cell stage. At 51-100-cell stage, 59.3% ± 4.2% of GFP signal remained in *pink-1(ok3538); allo-1(tm4756)* double mutant and 31.0% ± 5.1% in *allo-1(tm4756)* (***Figure 4A***). Our results indicate that, although the inactivation of *pink-1* in the *allo-1(tm4756)* mutant did not prevent the clearance of sperm-derived mitochondria, it increased their stability between the 10 and 100-cell stages.

**Figure 4.**
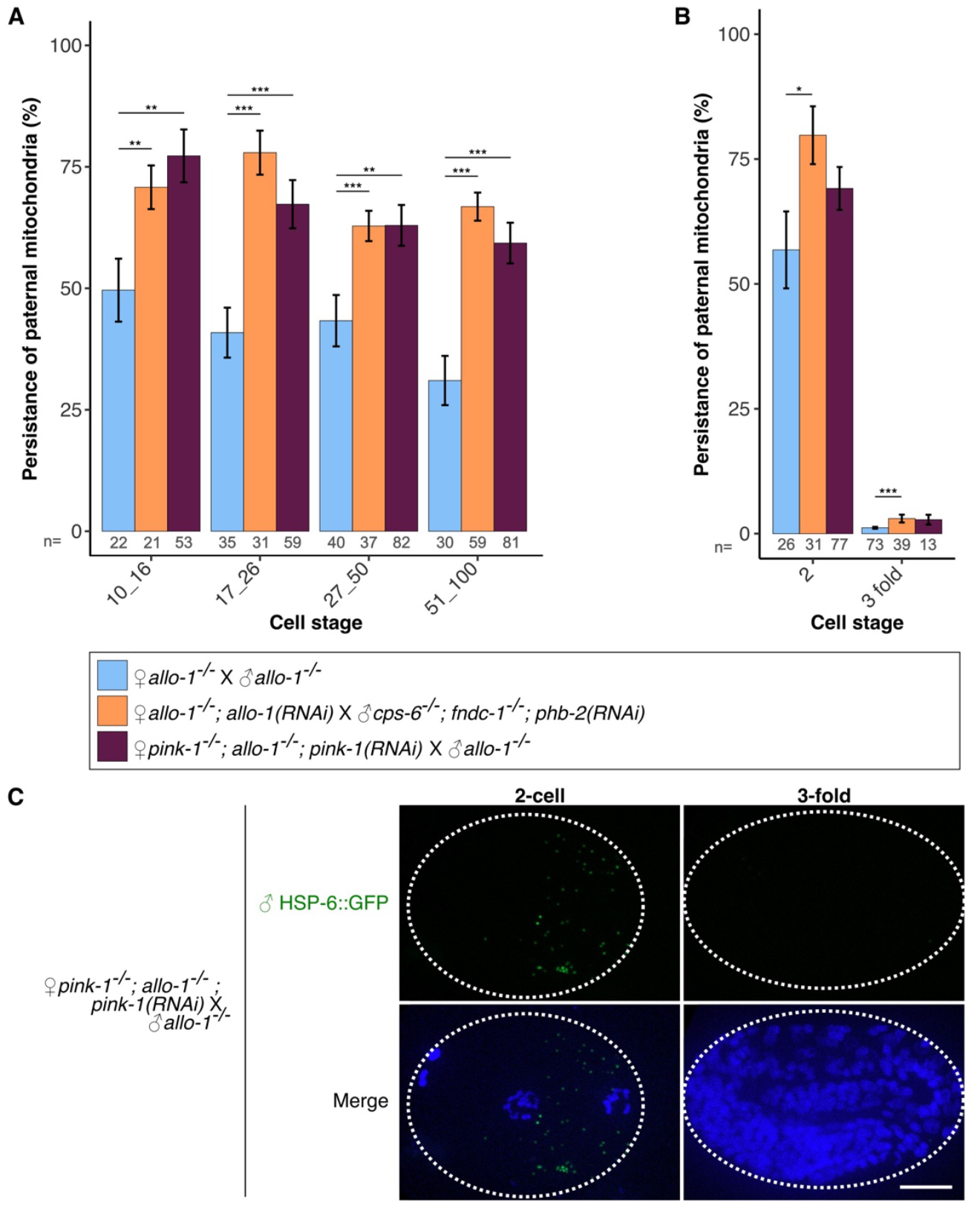
PINK-1 can target sperm-derived mitochondria in the absence of ALLO-1. **(A)** Quantification of the HSP-6::GFP labeled sperm-derived mitochondria (as % of the 1-cell stage) from the 10-cell to 100-cell stage and **(B)** in the late stage (3-fold stage). **(C)** Maximum intensity projections of confocal images of sperm-derived mitochondria (green) and nuclear DNA (blue) in 2-cell and late stage (3-fold) in embryos from the indicated crosses. Wilcoxon test p-values: *p<0.05; **p<0.01; ***p<0.001. Scale bar : 10µm.

## DISCUSSION

In most eukaryotes, mitochondria together with their mtDNA are maternally inherited. This uniparental maternal mode of heredity is very conserved in various vertebrate and invertebrate species. The *C. elegans* model system has been instrumental in understanding how sperm-derived mitochondria are eliminated after fertilization (Merlet et al., 2019). Several factors (ALLO-1, CPS-6, FNDC-1 and PHB-2) involved in sperm-derived mitochondria degradation have been identified, but how they interact together or not was unknown. Here, we provided new insights into the mechanisms driving sperm-derived mitochondria degradation in *C. elegans* embryos, specifically addressing how ALLO-1, CPS-6, FNDC-1, and PHB-2 cooperate or not in sperm mitochondria elimination.

### ALLO-1 and CPS-6 act synergistically in sperm mitochondrial degradation

One way to quantify sperm mitochondria after fertilization was to count the fluorescently labeled mitochondria. However, in early stages of development when sperm mitochondria are aggregated, counting individual mitochondria spots was almost impossible and resulted in a lot of variability. We developed a semi-autonomous method to quantify the mitochondrial HSP-6::GFP protein signal as a readout for sperm mitochondria quantification after fertilization which provides more accurate data. Using this method, we quantified sperm-derived mitochondria in double *cps-6(tm3222); allo-1(tm4756)* and in single *allo-1(tm4756)* and *cps-6(tm3222)* mutants. We observed that sperm-derived mitochondria remain longer in the double *cps-6(tm3222); allo-1(tm4756)* mutant compared to single mutants. Whereas we were able to reproduce published results from single *allo-1(tm4756)* mutant (Sato et al., 2018), we did not observe sperm mitochondria persistence on single *cps-6(tm3222)* mutant as published in Zhou et al. (Zhou et al., 2016). We also manually counted CMXRos labeled sperm mitochondria in *cps-6(tm3222)* mutants and did not find any mitochondria persistence (data not shown). The lack of effect we observed in single *cps-6(tm3222)* mutant suggested that, in *C. elegans*, CPS-6 is more dispensable on sperm-derived mitochondria elimination contrary to *Drosophila* where its homologous, the endonuclease endoG, seems to have a more central role (DeLuca and O’Farrell, 2012). Nevertheless, we observed an additive effect on stabilization of sperm-derived mitochondria on the double mutant *cps-6(tm3222); allo-1(tm4756)* which confirms the involvement of CPS-6 in the elimination of sperm-derived mitochondria. This result also suggested that ALLO-1 and CPS-6 act in pathways that are at least partially independent. The existence of several pathways acting in parallel was suggested by Zhou et al. which showed that 43% of sperm mitochondria are still surrounded by LGG-1 in *cps-6(tm3222)* mutant (Zhou et al., 2016). Our results provide evidence that at least two pathways co-exist, which act at least partially independently: one involving ALLO-1 and the other one CPS-6.

### Possible contribution of the PINK-1 pathway in *C. elegans*

We quantified sperm-derived mitochondria in double *pink-1(ok3538); allo-1(tm4756)* mutants. In early embryos, no additive effect was observed in this double mutant compared to single *allo-1(tm4756)* mutant. This finding corroborates previously published results, which demonstrated that degradation of sperm-derived mitochondria at early stages following fertilization was independent of PINK-1 (Sato et al., 2018). However, across all analyzed stages, we observed that sperm-derived mitochondria remain longer in double *pink-1(ok3538); allo-1(tm4756)* mutant than in single *allo-1(tm4756)* mutant between the 10-cell and 100-cell stages suggesting that PINK-1 could play a role in sperm-derived mitochondria elimination. Various previous studies have demonstrated that loss of mitochondrial membrane potential induces PINK-1 accumulation, which initiates the autophagic degradation of damaged mitochondria (Matsuda et al., 2010; Narendra et al., 2010; Vives-Bauza et al., 2010). Interestingly, we also reported that in the absence of allophagy, 16% of sperm-derived mitochondria lose their TMRE signal - a cationic mitochondrial fluorescent marker that stains mitochondria with intact membrane potential (Zorova et al., 2018) - at 20-24-cell stage (Rubio-Peña et al., 2021). Taken together, our results suggest that TMRE negative sperm mitochondria could be targeted to the autophagy machinery *via* PINK-1 but this hypothesis remains to be validated. Nevertheless, our results suggest that in an *allo-1* null sensitised background, the canonical mitophagy pathway mediated by PINK-1 may contribute to the degradation of at least a fraction of sperm-derived mitochondria.

### A nematode specific pathway is predominant for sperm mitochondria degradation after fertilization in *C. elegans*

*C. elegans* model system has been instrumental in deciphering the molecular mechanisms contributing to ensure a proper uniparental maternal mitochondrial heredity. Several proteins were identified independently: ALLO-1, CPS-6, FNDC-1 and PHB-2 (Lim et al., 2019; Sato et al., 2018; Wei et al., 2017; Zhou et al., 2016) but how they interact all together was unknown. Here, we analysed sperm-derived mitochondria fate in the embryo in the absence of all four proteins. Interestingly, we showed that in the quadruple mutant, sperm-derived mitochondria stabilization was the same as in the double mutant *cps-6(tm3222); allo-1(tm4756)* suggesting that all four proteins contributed to at least two pathways, one involving ALLO-1 and the other CPS-6. Furthermore, we showed that stabilization of sperm-derived mitochondria was not different in single *fndc-1(rny14)*, double *cps-6(tm3222); fndc-1(rny14)* and triple *cps-6(tm3222); fndc-1(rny14); phb-2(RNAi)* mutants suggesting that CPS-6, FNDC-1 and PHB-2 act in the same genetic pathway. Therefore, our results suggested that in *C. elegans* two pathways act in parallel, one involving ALLO-1 and the other one CPS-6, FNDC-1 and PHB-2 (***Figure 5***). Our quantifications of sperm-derived mitochondria in embryos also indicate that ALLO-1 pathway is the main pathway used in *C. elegans* to remove sperm mitochondria. Indeed, the inactivation of *allo-1* alone is enough to preserve 30 to 60% of sperm-derived mitochondria up to the 100-cell stage compared to other single mutants.

**Figure 5.**
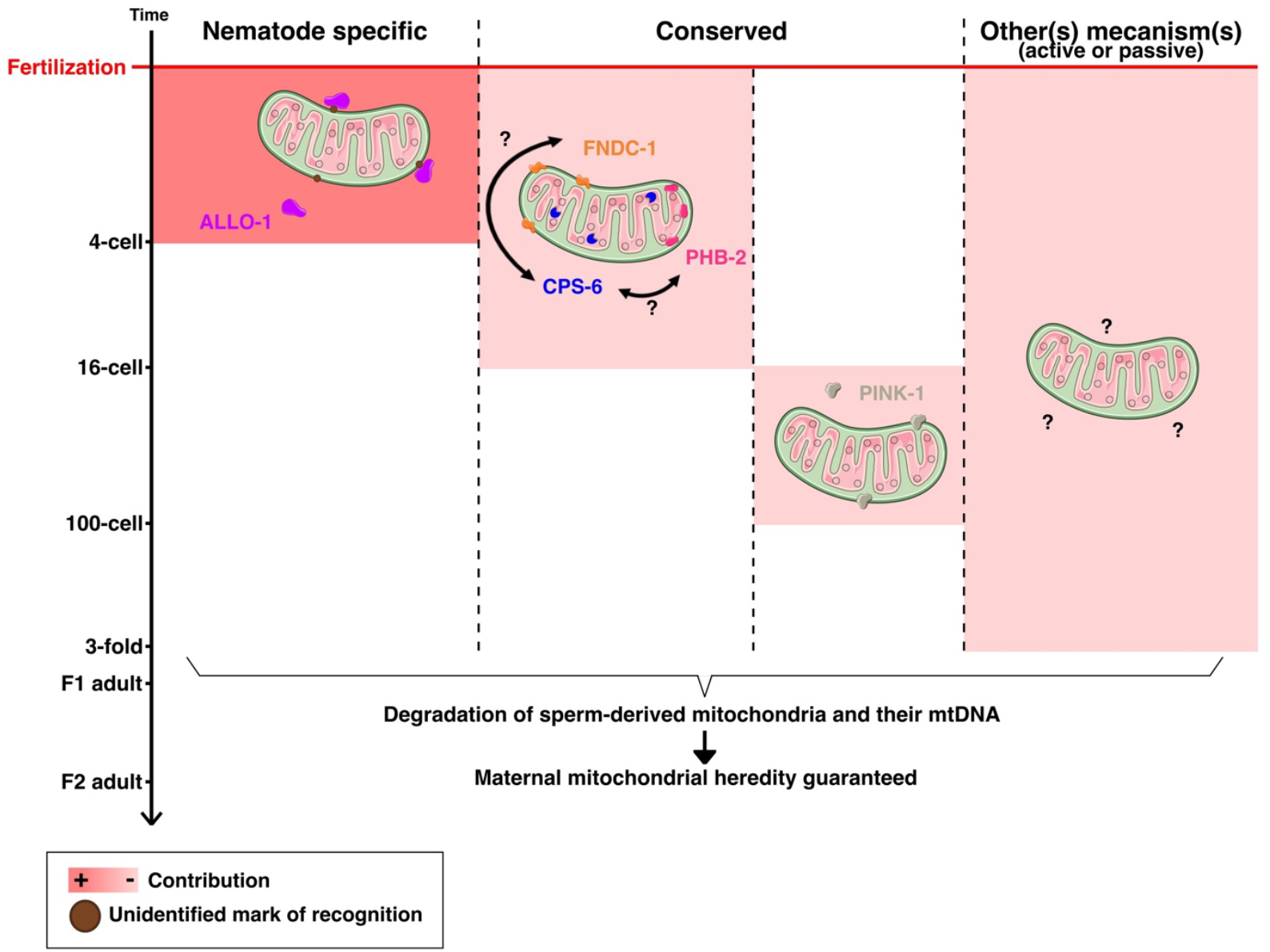
Putative model of the mechanisms involved in the degradation of sperm-derived mitochondria in *C. elegans*.

ALLO-1 is a nematode specific protein whereas CPS-6, FNDC-1 and PHB-2 are conserved proteins acting also in mitophagy of somatic cells. It seems that nematodes have developed a specific pathway to remove sperm mitochondria after fertilization but also conserved CPS-6, FNDC-1 and PHB-2 that might be part of an ancestral pathway. Altogether, those four proteins participate to ensure proper uniparental maternal mitochondrial heredity. Indeed, homologues of CPS-6 and PHB-2 have been found to participate in regulation of uniparental mitochondrial inheritance in other species like *Drosophila* and bivalves (DeLuca and O’Farrell, 2012; Wang et al., 2023). The role of FNDC-1 homologue in sperm-derived degradation in other species remains to be tested.

CPS-6, FNDC-1 and PHB-2 are all paternal factors; it will be interesting in the future to better understand when and how precisely they act during sperm development in *C. elegans*. Our analysis of the quadruple mutant showed that inactivation of those four factors is not sufficient to stabilize sperm-derived mitochondria. Indeed, in the quadruple mutant we observed that sperm-labeled mitochondria signal decreased in late embryos either when analyzed for GFP tagged HSP-6 protein or CMXRos staining. This suggests that other(s) mechanism(s) is(are) used in *C. elegans* to remove sperm-derived mitochondria and their mtDNA which remain(s) to be found.

In conclusion, our observations indicate that in *C. elegans* at least two main pathways act in parallel to ensure proper uniparental maternal mitochondrial heredity: a minor one involving the conserved proteins CPS-6, FNDC-1 and PHB-2 and a major worm specific pathway, involving the autophagy receptor ALLO-1.

## MATERIAL AND METHODS

### Key resource table

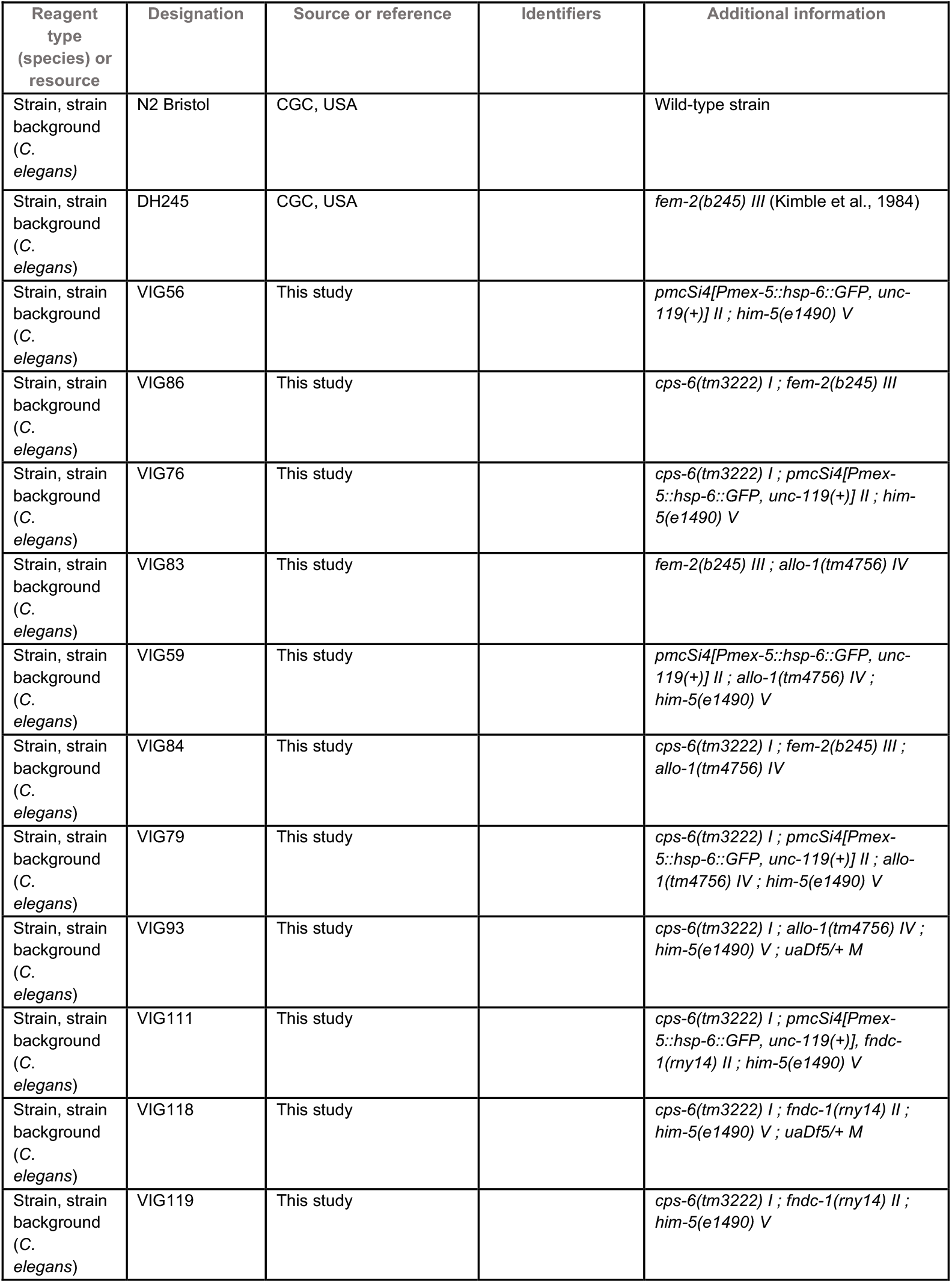
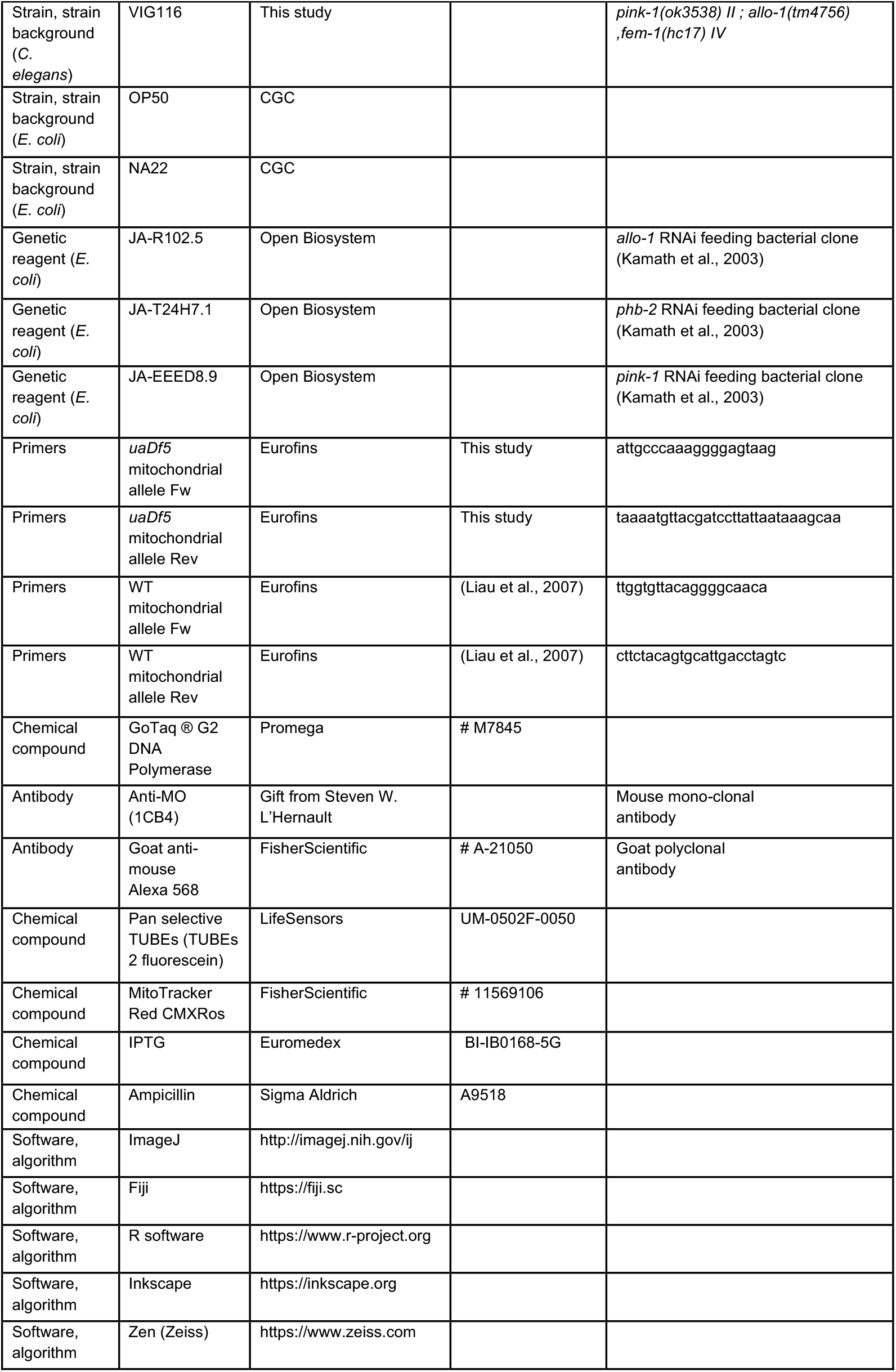

#### *C. elegans* strains, maintenance and RNAi

Strains were cultured and maintained under standard procedures (Brenner, 1974). *C. elegans* and bacterial strains used in this study are listed in the key resource table. RNAi was performed by the feeding method as described (Kamath et al., 2001) on NGM plates containing 100 µg/mL ampicillin and IPTG (2 mM). For *phb-2(RNAi)*, young adult males were maintained on RNAi plates for 24h at 20°C. For *allo-1(RNAi)* and *pink-1(RNAi)*, animals were grown from L1 larvae for 48h at 25°C. Then, males and feminised worms were transferred to new RNAi plates at 20°C to cross. For negative control, animals were fed on HT115 bacteria containing the empty L4440 plasmid. RNAi clones were obtained from the Ahringer library (Open Source BioScience).

#### Mating experiments

Males were obtained from synchronized L1 worms, plated on 10 cm NGM plates seeded with *E. coli* strain NA22 for 72h at 20°C as described (Cailloce et al., 2023b). For all crosses we used feminized hermaphrodites from 10-20 *fem* adult hermaphrodites placed at 25°C for 48h to 72h. For the crosses, 40 young feminized worms and 120 young males of the genotype of interest were transferred to a new plate with a small spot of OP50 or HTT115 (for RNAi experiments) bacteria and allowed to mate for 24h at 20°C.

#### Dissection and fixation

To obtain embryos at late stages of development, hermaphrodites were placed in 50 µL of meiosis medium (60 % Leibovitz’s L-15 Medium no phenol red, 20 % FBS, 0.5 mg/ml Inulin, 25 mM HEPES) in a humid chamber at 25°C for 2 to 7 hours depending on the desired stage. For the imaging of fixed embryos, around 15 gravid hermaphrodites were dissected in a 10 μL drop of M9 medium (Na2HPO4 33.9 g/L; KH2PO4 15 g/L; NH4Cl 5 g/L; NaCl 2.5 g/L) on a microscope slide pre-coated with a “poly-lysine solution” (gelatin 2%, chromium III 0.2 g/L, sodium azide 1 mM, poly-L-lysine 1 g/L). A 10×10 mm coverslip is then placed over the sample before snap-freezing the slide on dry ice. The coverslip is removed by “popping” them with a scalpel. Embryos were fixed in dehydrated methanol at −20°C for 30 minutes. Slides are rinsed for 10 minutes in PBS-T (Phosphate Buffered Salin with 0.1% Tween-20) at room temperature, then mounted with 8 μL of Vectashield-DAPI (Vector Lab). Samples are sealed with molten VALAP solution (a 1:1:1 mixture of petroleum jelly, lanolin and paraffin wax).

#### CMXRos staining

To stain male mitochondria, young males were transferred on NGM plates containing Red CMXRos (1 µg/mL, FisherScientific) and incubated overnight in the dark. Stained males were then transferred with feminized hermaphrodites for crosses as described above.

#### Immunofluorescence

After methanol fixation, embryos were washed with PBS-T and blocked in PBS-T-milk (5% milk). Embryos were washed in PBS-T and incubated in a wet chamber protected from light for 2 hours at room temperature in PBS-T-milk containing fluorescein labeled TUBEs (1/1000). For MOs staining, 1CB4 anti-MO (1/2000) was added together with TUBEs. Then slides were washed in PBS-T and incubated for 1h at room temperature with anti-mouse Alexa 568 (1/800) (Cailloce et al., 2023a). Slides were then washed 3 times in PBS-T, once in PBS and mounted as described above.

#### Worms’ and embryos lysis

Worms were put in 10 µL of lysis buffer (50 mM KCl, 10 mM Tris, 20 mM MgCl2, 0.45% NP40, 0.45% Tween-20, 0.01% gelatin) containing 1 mg/ml proteinase K, flash frozen in liquid nitrogen, incubated at 60°C for 1h, followed by 10 min at 95°C. For embryo lysis, 20 embryos were washed three times in M9 using a mouth pipet, once in M9+Triton 0,1% and transferred in 10 µL of lysis solution and lysed.

#### mtDNA PCR conditions

PCR reactions were performed with the GoTaq G2 DNA polymerase (Promega) following manufacturer’s instructions. PCR primers used in this study are listed in the key resource table. For *uaDf5* mtDNA amplification, the PCR was performed as follows: 2 min at 95°C, 35 cycles of 95°C for 30 seconds, 53°C for 30 seconds and 72°C for 1 min, followed by 5 min at 72°C. For WT mtDNA amplification, the PCR conditions were: 2 min at 95°C, 30 cycles of 95°C for 30 s, 51°C for 30 seconds and 72°C for 40 s, followed by 5 min at 72°C.

#### Confocal imaging

Embryos were imaged using a spinning disk confocal microscope (Cell Observer Zeiss) equipped with a rotating-disk confocal head (Yokogawa CSU-X1), a back-illuminated CCD camera (Evolve 512 EM) and a 100X 1.46NA immersion objective. For each embryo, the whole embryonic volume was imaged by acquiring Z-stacks of 76 images at 0.4 μm intervals. Each embryo was exposed consecutively to 568 nm, 488 nm and 405 nm lasers, for CMXRos or MOs staining, GFP and DAPI respectively. A single transmisted white light acquisition was performed on the median plane using Differential Interference Contrast (DIC) optics.

#### Image analysis and quantification

For each embryo, the intensity of fluorescence from sperm mitochondria was measured with Fiji software (Schindelin et al., 2012) (https://fiji.sc). Orthogonal projection of the summed intensities was performed in all fluorescence channels from the 76 stacks. The DAPI-summed image was used to determine the stage of embryonic development by counting the number of nuclei. The green channel was used to measure the total fluorescence intensity of sperm mitochondria per embryo in the region of interest (ROI) defined from the DIC acquisition to contain the whole embryo. The threshold was chosen to subtract background fluorescence. For MOs and CMXRos-labeled mitochondria quantifications, particles were counted over the entire volume of the embryo.

#### Statistical analysis

All statistical analyses and graphs were performed using R software (https://www.r-project.org). A Wilcoxon statistical test was performed on the data except for Figure S1A where a Student test was performed. Results are presented as mean ± standard deviation from the mean (SEM). The significance level, or p-value, is defined as less than or equal to 0.05 (p ≤ 0.05).

#### Software

Figures were assembled using Inkscape software (https://inkscape.org/). Icons and images were found on Bioicons (https://bioicons.com/) and NIAID NIH BIOART Source (https://bioart.niaid.nih.gov/).

## ACKNOWLEDGEMENTS

We are very grateful to Ritha Zamy for valuable technical assistance. We are also very grateful to Professor Steven W. L’Hernault and Elizabeth Gleason for giving us the 1CB4 antibody and sharing their protocole. We thank Dr. Lionel Pintard and Pr. Clément Carré for critical reading of the manuscript. Some strains were provided by the CGC, which is funded by NIH Office of Research Infrastructure Programs (P40 OD010440). This work was supported by the Agence Nationale de la Recherche, France (ANR 12-BSV2-0018–01 and ANR-22-CE13–0042 to VG).

---

**Figure S1.**
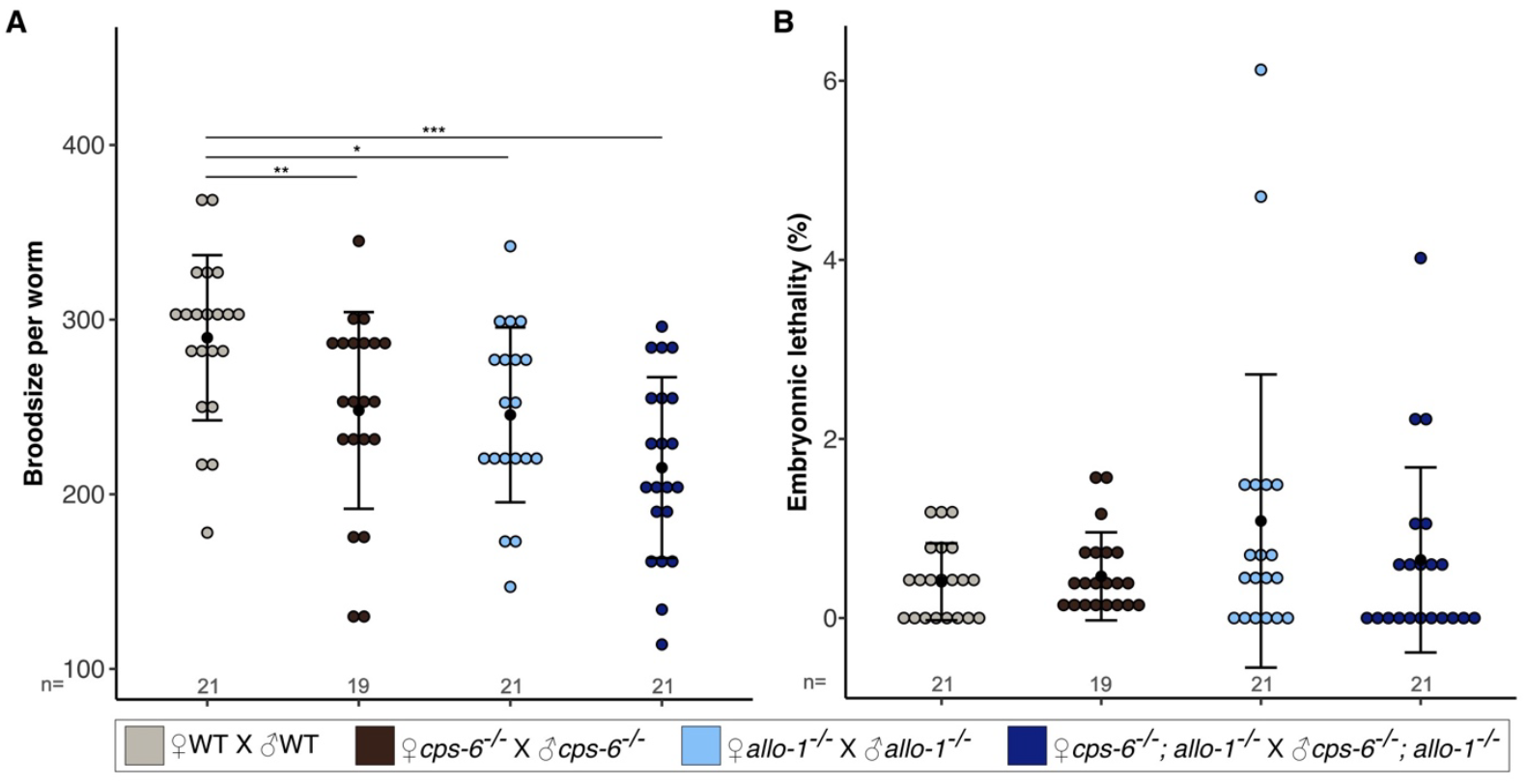
No deleterious effects on *allo-1(tm4756)* and *cps-6(tm3222)* simple and double mutants embryos. **(A)** Quantification of the progeny per worm and **(B)** quantification of the embryonic lethality. Student test (A) and Wilcoxon (B) test p-values: *p<0.05; **p<0.01; ***p<0.001.

## REFERENCES

1. Al Rawi S, Louvet-Vallée S, Djeddi A, Sachse M, Culetto E, Hajjar C, Boyd L, Legouis R, Galy V. 2011. Postfertilization Autophagy of Sperm Organelles Prevents Paternal Mitochondrial DNA Transmission. Science (1979) 334:1144–1147. doi:10.1126/science.1211878

2. Birky CW. 2001. The Inheritance of Genes in Mitochondria and Chloroplasts: Laws, Mechanisms, and Models. Annu Rev Genet 35:125–148. doi:10.1146/annurev.genet.35.102401.090231

3. Brenner S. 1974. The genetics of Caenorhabditis elegans. Genetics 77:71–94. doi:10.1093/genetics/77.1.71

4. Cailloce J, Husson F, Galy V, Merlet J. 2023a. An antibody free approach to probe the presence of poly-ubiquitin chains on C. elegans sperm derived organelles after fertilization. MicroPubl Biol 2023. doi:10.17912/micropub.biology.000972

5. Cailloce J, Husson F, Zablocki A, Galy V, Merlet J. 2023b. Fast and easy method to culture and obtain large populations of male nematodes. MethodsX 11:102293. doi:10.1016/j.mex.2023.102293

6. DeLuca SZ, O’Farrell PH. 2012. Barriers to Male Transmission of Mitochondrial DNA in Sperm Development. Dev Cell 22:660–668. doi:10.1016/j.devcel.2011.12.021

7. Kamath RS, Fraser AG, Dong Y, Poulin G, Durbin R, Gotta M, Kanapin A, Le Bot N, Moreno S, Sohrmann M, Welchman DP, Zipperlen P, Ahringer J. 2003. Systematic functional analysis of the Caenorhabditis elegans genome using RNAi. Nature 421:231–237. doi:10.1038/nature01278

8. Kamath RS, Martinez-Campos M, Zipperlen P, Fraser AG, Ahringer J. 2001. Effectiveness of specific RNA-mediated interference through ingested double-stranded RNA in Caenorhabditis elegans. Genome Biol 2:1–10. doi:10.1186/gb-2000-2-1-research0002

9. Kimble J, Edgar L, Hirsh D. 1984. Specification of male development in Caenorhabditis elegans: The fem genes. Dev Biol 105:234–239. doi:10.1016/0012-1606(84)90279-3

10. Liau W-S, Gonzalez-Serricchio AS, Deshommes C, Chin K, LaMunyon CW. 2007. A persistent mitochondrial deletion reduces fitness and sperm performance in heteroplasmic populations of C. elegans. BMC Genet 8:8. doi:10.1186/1471-2156-8-8

11. Lim Y, Rubio-Peña K, Sobraske PJ, Molina PA, Brookes PS, Galy V, Nehrke K. 2019. Fndc-1 contributes to paternal mitochondria elimination in C. elegans. Dev Biol 454:15–20. doi:10.1016/j.ydbio.2019.06.016

12. Matsuda N, Sato S, Shiba K, Okatsu K, Saisho K, Gautier CA, Sou YS, Saiki S, Kawajiri S, Sato F, Kimura M, Komatsu M, Hattori N, Tanaka K. 2010. PINK1 stabilized by mitochondrial depolarization recruits Parkin to damaged mitochondria and activates latent Parkin for mitophagy. Journal of Cell Biology 189. doi:10.1083/jcb.200910140

13. Merlet J, Rubio-Peña K, Al Rawi S, Galy V. 2019. Autophagosomal Sperm Organelle Clearance and mtDNA Inheritance in C. elegansAdvances in Anatomy Embryology and Cell Biology. pp. 1–23. doi:10.1007/102_2018_1

14. Narendra DP, Jin SM, Tanaka A, Suen DF, Gautier CA, Shen J, Cookson MR, Youle RJ. 2010. PINK1 is selectively stabilized on impaired mitochondria to activate Parkin. PLoS Biol 8. doi:10.1371/journal.pbio.1000298

15. Parrish J, Li L, Klotz K, Ledwich D, Wang X, Xue D. 2001. Mitochondrial endonuclease G is important for apoptosis in C. elegans. Nature 412. doi:10.1038/35083608

16. Rubio-Peña K, Al Rawi S, Husson F, Lam F, Merlet J, Galy V. 2021. Mitophagy of polarized sperm-derived mitochondria after fertilization. iScience 24:102029. doi:10.1016/j.isci.2020.102029

17. Sato M, Sato K. 2011. Degradation of Paternal Mitochondria by Fertilization-Triggered Autophagy in C. elegans Embryos. Science (1979) 334:1141–1144. doi:10.1126/science.1210333

18. Sato M, Sato Katsuya, Tomura K, Kosako H, Sato Ken. 2018. The autophagy receptor ALLO-1 and the IKKE-1 kinase control clearance of paternal mitochondria in Caenorhabditis elegans. Nat Cell Biol 20:81–91. doi:10.1038/s41556-017-0008-9

19. Schindelin J, Arganda-Carreras I, Frise E, Kaynig V, Longair M, Pietzsch T, Preibisch S, Rueden C, Saalfeld S, Schmid B, Tinevez J-Y, White DJ, Hartenstein V, Eliceiri K, Tomancak P, Cardona A. 2012. Fiji: an open-source platform for biological-image analysis. Nat Methods 9:676–682. doi:10.1038/nmeth.2019

20. Vives-Bauza C, Zhou C, Huang Y, Cui M, De Vries RLA, Kim J, May J, Tocilescu MA, Liu W, Ko HS, Magrané J, Moore DJ, Dawson VL, Grailhe R, Dawson TM, Li C, Tieu K, Przedborski S. 2010. PINK1-dependent recruitment of Parkin to mitochondria in mitophagy. Proc Natl Acad Sci U S A 107. doi:10.1073/pnas.0911187107

21. Wang Y, Zhu X, Gu Y, Liu Z, Mao Y, Liu X, Bai Z, Wang G, Li J. 2023. Study on the Role of Mitophagy Receptor PHB2 in Doubly Uniparental Inheritance of Hyriopsis cumingii. Marine Biotechnology 25. doi:10.1007/s10126-023-10240-5

22. Wei Y, Chiang W-C, Sumpter R, Mishra P, Levine B. 2017. Prohibitin 2 Is an Inner Mitochondrial Membrane Mitophagy Receptor. Cell 168:224-238.e10. doi:10.1016/j.cell.2016.11.042

23. Zhou Q, Li H, Li H, Nakagawa A, Lin JLJ, Lee E-S, Harry BL, Skeen-Gaar RR, Suehiro Y, William D, Mitani S, Yuan HS, Kang B-H, Xue D. 2016. Mitochondrial endonuclease G mediates breakdown of paternal mitochondria upon fertilization. Science (1979) 353:394–399. doi:10.1126/science.aaf4777

24. Zorova LD, Popkov VA, Plotnikov EJ, Silachev DN, Pevzner IB, Jankauskas SS, Zorov SD, Babenko VA, Zorov DB. 2018. Functional Significance of the Mitochondrial Membrane Potential. Biochem (Mosc) Suppl Ser A Membr Cell Biol. doi:10.1134/S1990747818010129

